# Evolutionary stability of topologically associating domains is associated with conserved gene regulation

**DOI:** 10.1101/231431

**Authors:** Jan Krefting, Miguel A. Andrade-Navarro, Jonas Ibn-Salem

## Abstract

**Background:** The human genome is highly organized in the three-dimensional nucleus. Chromosomes fold locally into topologically associating domains (TADs) defined by increased intra-domain chromatin contacts. TADs contribute to gene regulation by restricting chromatin interactions of regulatory sequences, such as enhancers, with their target genes. Disruption of TADs can result in altered gene expression and is associated to genetic diseases and cancers. However, it is not clear to which extent TAD regions are conserved in evolution and whether disruption of TADs by evolutionary rearrangements can alter gene expression.

**Results:** Here, we hypothesize that TADs represent essential functional units of genomes, which are selected against rearrangements during evolution. We investigate this using whole-genome alignments to identify evolutionary rearrangement breakpoints of different vertebrate species. Rearrangement breakpoints are strongly enriched at TAD boundaries and depleted within TADs across species. Furthermore, using gene expression data across many tissues in mouse and human, we show that genes within TADs have more conserved expression patterns. Disruption of TADs by evolutionary rearrangements is associated with changes in gene expression profiles, consistent with a functional role of TADs in gene expression regulation.

**Conclusions:** Together, these results indicate that TADs are conserved building blocks of genomes with regulatory functions that are often reshuffled as a whole instead of being disrupted by rearrangements.

## Background

The three-dimensional structure of eukaryotic genomes is organized in many hierarchical levels [1]. The development of high-throughput experiments to measure pairwise chromatin-chromatin interactions, such as Hi-C [2] enabled the identification of genomic domains of several hundred kilo-bases with increased self-interaction frequencies, described as topologically associating domains (TADs) [3–5]. Loci within TADs contact each other more frequently and TAD boundaries insulate interactions of loci in different TADs. TADs have also been shown to be important for gene regulation by restricting the interaction of cell-type specific enhancers with their target genes [4,6,7]. Several studies associated disruption of TADs to ectopic regulation of important developmental genes leading to genetic diseases [8–10]. These properties of TADs suggested that they are functional genomic units of gene regulation.

Interestingly, TADs are largely stable across cell-types [3,11] and during differentiation [12]. Moreover, while TADs were initially described for mammalian genomes, a similar domain organization was found in the genomes of non-mammalian species such as *Drosophila* [5], zebrafish [13] *Caenorhabditis elegans* [14] and yeast [15,16]. Evolutionary conservation of TADs together with their spatio-temporal stability within organisms, would collectively imply that TADs are robust structures.

This motivated the first studies comparing TAD structures across different species, which indeed suggested that individual TAD boundaries are largely conserved along evolution. More than 54% of TAD boundaries in human cells occur at homologous positions in mouse genomes [3]. Similarly, 45% of contact domains called in mouse B-lymphoblasts were also identified at homologous regions in human lymphoblastoid cells [11]. A single TAD boundary at the Six gene loci could be traced back in evolution to the origin of deuterostomes [13]. However, these analyses focused only on the subset of syntenic regions that can be mapped uniquely between genomes and do not investigate systematically if TAD regions as a whole might be stable or disrupted by rearrangements during evolution.

A more recent study provided Hi-C interaction maps of liver cells for four mammalian genomes [17]. Interestingly, they described three examples of rearrangements between mouse and dog, which all occurred at TAD boundaries. However, the rearrangements were identified by ortholog gene adjacencies, which might be biased by gene density. Furthermore, they did not report the total number of rearrangements identified, leaving the question open of how many TADs are actually conserved between organisms. It remains unclear to which extent TADs are selected against disruptions during evolution [18]. All these studies underline the need to make a systematic study to verify if and how TAD regions as a whole might be stable or disrupted by rearrangements during evolution.

To address this issue we used whole-genome alignment data to analyze systematically whether TADs represent conserved genomic structures that are rather reshuffled as a whole than disrupted by rearrangements during evolution. Furthermore, we used gene expression data from many tissues in human and mouse to associate disruptions of TADs by evolutionary rearrangements to changes in gene expression.

## Results

### Identification of evolutionary rearrangement breakpoints from whole-genome alignments

To analyze the stability of TADs in evolution, we first identified evolutionary rearrangements by using whole-genome alignment data from the UCSC Genome Browser [19,20] to compare the human genome to 12 other species. These species where selected to have genome assemblies of good quality and to span several hundred million years of evolution. They range from chimpanzee to zebrafish (Figure 1). The whole-genome data consists of consecutive alignment blocks that are chained and hierarchically ordered into so-called net files as fills [19]. To overcome alignment artifacts and smaller local variations between genomes we only considered top-level fills or non-syntenic fills and additionally applied a size threshold to use only fills that are larger than 10 kb, 100 kb, or 1000 kb, respectively. Start and end coordinates of such fills represent borders of syntenic regions and were extracted as rearrangement breakpoints for further analysis (see Methods for details).

**Fig. 1.**
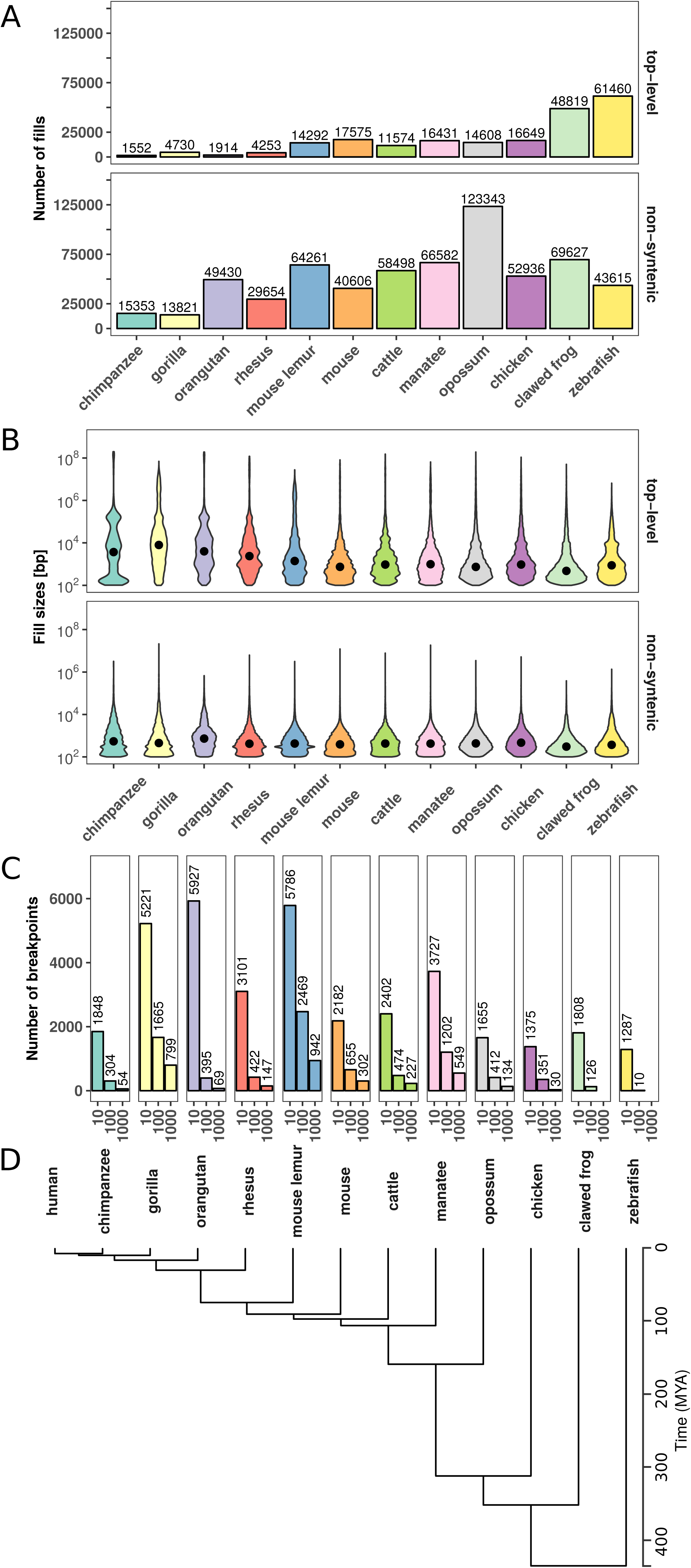
Number and size distributions of fill sizes of whole-genome alignments between human and 12 other species. **(A)** Number of syntenic alignment blocks (fills) between human (hg38) and 12 other species. Top-level fills are the largest and highest scoring chains and occur at the top level in the hierarchy in net files (top panel). Non-syn fills map to different chromosomes as their parent fills in the net files (bottom panel). **(B)** Size distribution of top-level (top panel) and non-syntenic (bottom panel) fills as violin plot. **(C)** Number of identified rearrangement breakpoints between human and 12 other species. Breakpoints are borders of top-level or non-syn fills that are larger or equal than a given size threshold (x-axis). **(D)** Phylogenetic tree with estimated divergence times according to http://timetree.org/.

First, we analyzed the number and size distributions of top-level and non-syntenic fills between human and other species (Fig. 1). As expected, closely related species such as chimpanzee and gorilla have in general fewer fills but larger fill sizes (mean length ≥1 kb), whereas species which are more distant to human, such as chicken and zebrafish, tend to have more but smaller fills (mean length ≤ 1 kb, Fig. 1A,B). However, we also observe many small non-syntenic fills in closely related species, likely arising from transposon insertions [22]. As a consequence of the number of fills and size distributions, we identify different breakpoint numbers depending on species and size threshold applied. For example, the whole-genome alignment between human and mouse results in 2182, 655, and 302 rearrangement breakpoints for size thresholds, 10 kb, 100 kb, and 1000 kb, respectively (Fig. 1C). Together, the number and size distributions of syntenic regions reflect the evolutionary divergence time from human and allow us to identify thousands of evolutionary rearrangement breakpoints for enrichment analysis at TADs.

### Rearrangement breakpoints are enriched at TAD boundaries

Next, we analyzed how the identified rearrangement breakpoints are distributed in the human genome with respect to TADs. We obtained 3,062 TADs identified in human embryonic stem cells (hESC) [3] and 9,274 contact domains from high-resolution *in situ* Hi-C in human B-lymphoblastoid cells (GM12878) [22]. To calculate the number of breakpoints around TADs, we enlarged each TAD region by +/-50% of its size and divided the region in 20 equal sized bins. For each bin we computed the number of overlapping rearrangement breakpoints. This results in a size-normalized distribution of rearrangement breakpoints along TAD regions. First, we analyzed the distribution of breakpoints at different size thresholds between human and mouse at hESC TADs (Fig. 2A). Rearrangement breakpoints are clearly enriched at TAD boundaries and depleted within TAD regions. Notably, this enrichment is observed for all size thresholds applied in the identification of rearrangement breakpoints. Next, we also analyzed the breakpoints from chimpanzee, cattle, opossum, and zebrafish (Fig. 2B) at the 10 kb size threshold. Interestingly, we observed for all species a clear enrichment of breakpoints at TAD boundaries and depletion within TAD regions. To quantify this enrichment, we simulated an expected background distribution of breakpoints by placing each breakpoint 100 times at a random position of the respective chromosome. We than calculated the fraction of observed and expected breakpoints that are closer than 40 kb to a TAD boundary. For all size thresholds and analyzed species, we computed the log-fold-ratio of actual breakpoints over random breakpoints at domain boundaries (Fig. 2C). For virtually all species and size thresholds analyzed, we found breakpoints significantly enriched at boundaries of TADs and contact domains (Fig. 2C, S1). Depletion was only observed for some combinations of species and size thresholds which have only very few breakpoints (see Fig. 1C). Furthermore, we compared the distance of each breakpoint to the closest TAD boundary and observed nearly always significantly shorter distances for actual breakpoints compared to random controls (Fig. S2). Overall, the enrichment was stronger for TADs in hESC compared to the contact domains in GM12878. However, these differences were likely due to different sizes of TADs and contact domains and the nested structure of contact domains, which overlap each other [22]. Rearrangements between human and both closely and distantly related species are highly enriched at TAD boundaries and depleted within TADs. These results show (i) that rearrangements are not randomly distributed in the genome, in agreement with [23], and (ii) strong conservation of TAD regions over large evolutionary time scales, indicating selective pressure against disruption of TADs, presumably because of their functional role in gene expression regulation.

**Fig. 2.**
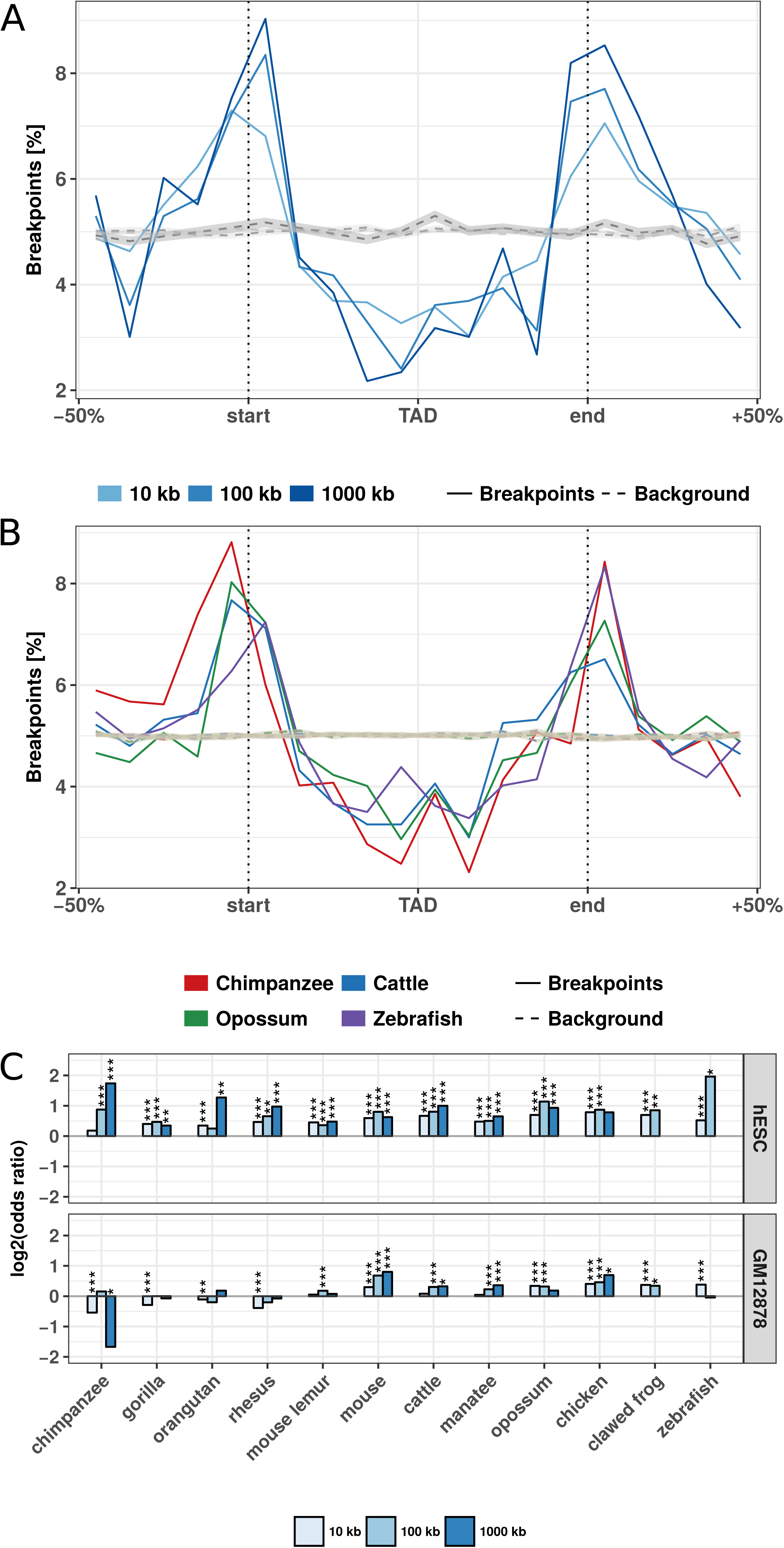
Evolutionary rearrangements are enriched at TAD boundaries. **(A)** Distribution of evolutionary rearrangement breakpoints between human and mouse around hESC TADs. Each TAD and 50% of its adjacent sequence was subdivided into 20 bins of equal size, the breakpoints were assigned to the bins and their number summed up over the corresponding bins in all TADs. Blue color scale represents breakpoints from different fill-size thresholds. Dotted lines in gray show simulated background controls of randomly placed breakpoints. **(B)** Distribution of rearrangement breakpoints between human and: chimpanzee, cattle, opossum, and zebrafish, at 10 kb size threshold around hESC TADs. Dotted lines in gray show simulated background controls of randomly placed breakpoints. **(C)** Enrichment of breakpoints at TAD boundaries as log-odds-ratio between actual breakpoints at TAD boundaries and randomly placed breakpoints. Enrichment is shown for three different fill size thresholds (blue color scale) and TADs in hESC from [3] (top) and contact domains in human GM12878 cells from [22] (bottom), respectively. Asterisks indicate significance of the enrichment using Fisher's exact test (*p <= 0.05; **p <= 0.01; ***p <= 0.001).

### Clusters of conserved non-coding elements are depleted for rearrangement breakpoints

Another interesting feature that can be extracted from whole-genome alignments are highly conserved non-coding elements (CNEs) [24]. CNEs are defined as non-protein-coding sequences of at least 50 bp with over 70% sequence identity between distantly related species such as human and chicken [24]. In the human genome, CNEs cluster around developmental genes in so-called genomic regulatory blocks (GRBs) [25]. It has been shown recently that many GRBs coincide with TADs in human and *Drosophila* genomes [26]. Therefore, we asked whether evolutionary breakpoints are also enriched at boundaries of GRBs. This would support the idea of a conserved regulatory environment around important developmental genes. Indeed we saw a strong enrichment around GRBs (Fig. 3A). This is consistent with previous studies in *Drosophila* and Fish where CNE arrays often correspond to syntenic blocks [27,28]. Next, we subdivided TADs according to their overlap with GRBs in GRB-TADs (> 80% overlap) and non-GRB-TADs (< 20% overlap) as in the original study [26]. As expected, we observed a higher accumulation of breakpoints at boundaries and stronger depletion within TADs for GRB-TADs compared to non-GRB-TADs (Fig. 3B). However, also the non-GRB-TADs, that have less than 20% overlap with GRBs, are enriched for rearrangements at TAD boundaries. This indicates that not only TADs overlapping GRBs are evolutionary conserved. In summary, we show that human TADs overlapping clusters of non-coding conserved elements are strongly depleted for rearrangements, likely due to strong selective pressure on the conserved regulatory environment around important developmental genes.

**Fig. 3.**
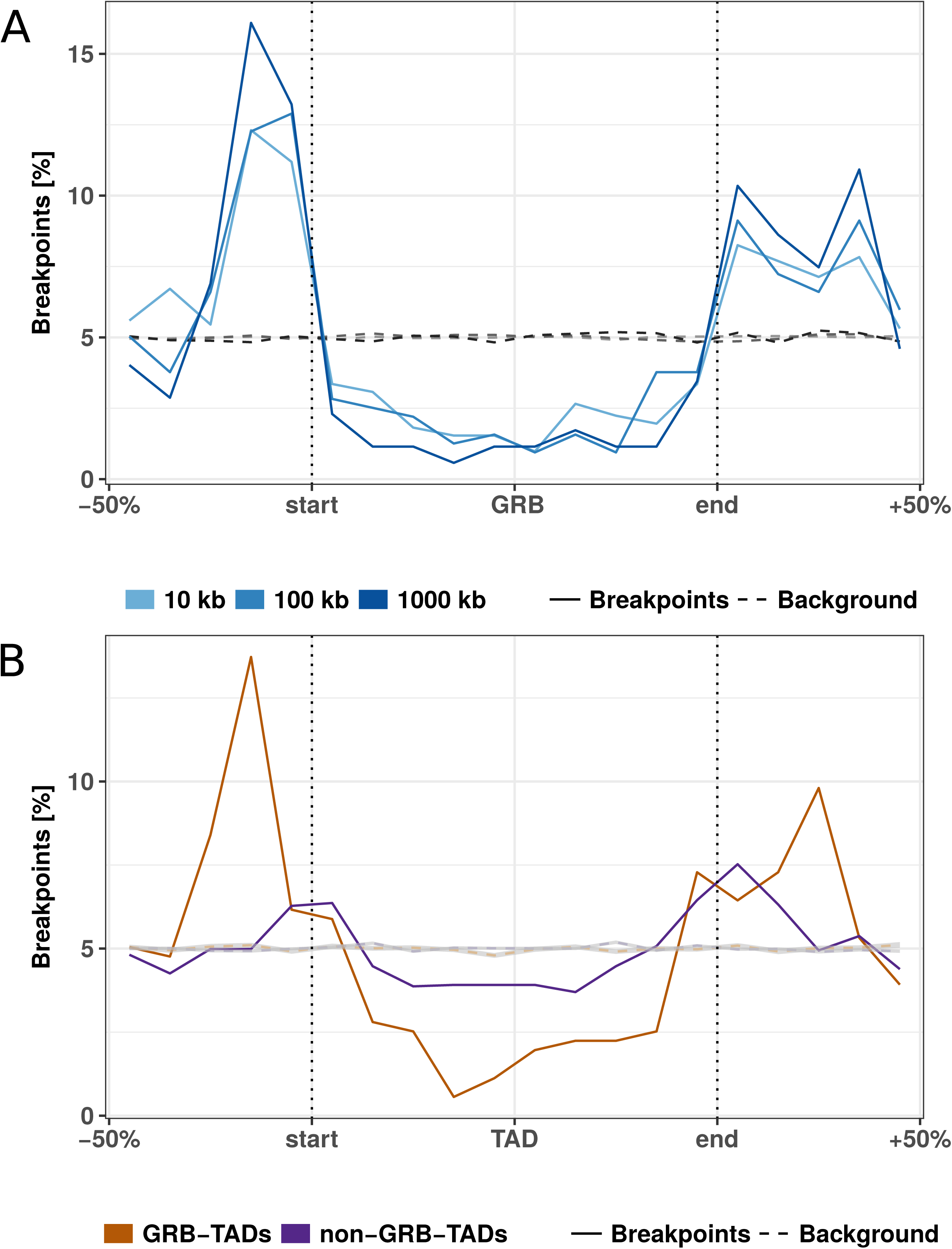
Rearrangement breakpoint distribution around GRBs and GRB-TADs. **(A)** Rearrangement breakpoints between mouse and human around 816 GRBs. **(B)** Breakpoint distribution around GRB-TADs and non-GRB-TADs. GRB-TADs are defined as TADs overlapping more than 80% with GRBs and non-GRB-TADs have less than 20% overlap with GRBs. Breakpoints using a 10 kb fill size threshold are shown.

### Rearranged TADs are associated with divergent gene expression between species

The enrichment of rearrangement breakpoints at TAD boundaries indicates that TADs are stable across large evolutionary time scales. However, the reason for this strong conservation of TAD regions is unclear. A mechanistic explanation could be that certain chromatin features at TAD boundaries promote or prevent DNA double strand breaks (DSBs) [23,29]. Alternatively, selective pressure might act against the disruption of TADs due to their functional importance, for example in developmental gene regulation [30,31]. TADs constitute a structural framework determining possible interactions between promoters and cis-regulatory sequences while prohibiting the influence of other sequences [6,9]. TAD disruption would prevent formerly established contacts. Rearrangements of TADs might also enable the recruitment of new cis-regulatory sequences which would alter the expression patterns of genes in rearranged TADs [9,32]. Because of these detrimental effects, rearranged TADs should largely be eliminated by purifying selection. However, rearrangement of TADs could also enable the expression of genes in a new context and be selected if conferring an advantage. Therefore, we hypothesized that genes within conserved TADs might have a more stable gene expression pattern across tissues, whereas genes in rearranged TADs between two species might have a more divergent expression between species.

To test this, we analyzed the conservation of gene expression of ortholog genes between human and mouse across 19 matched tissues from the FANTOM5 project (Table. S1) [33]. If a human gene and its mouse ortholog have high correlation across matching tissues, they are likely to have the same regulation and eventually similar functions. Conversely, low correlation of expression across tissues can indicate functional divergence during evolution, potentially due to altered gene regulation.

First, we separated human genes according to their location within TADs or outside of TADs. From 12,696 human genes with expression data and a unique one-to-one ortholog in mouse (Table S2), 1,525 have a transcription start site (TSS) located outside hESC TADs and 11,171 within. Next, we computed for each gene its expression correlation with mouse orthologs across 19 matching tissues. Genes within TADs have significantly higher expression correlation with their mouse ortholog (median R = 0,340) compared to genes outside TADs (mean R = 0,308, p = 0.0015, Fig. 4A). This indicates higher conservation of gene regulation in TADs and is consistent with the observation of housekeeping genes at TAD boundaries [3] and the role of TADs in providing conserved regulatory environments for gene regulation [27,35].

**Fig. 4.**
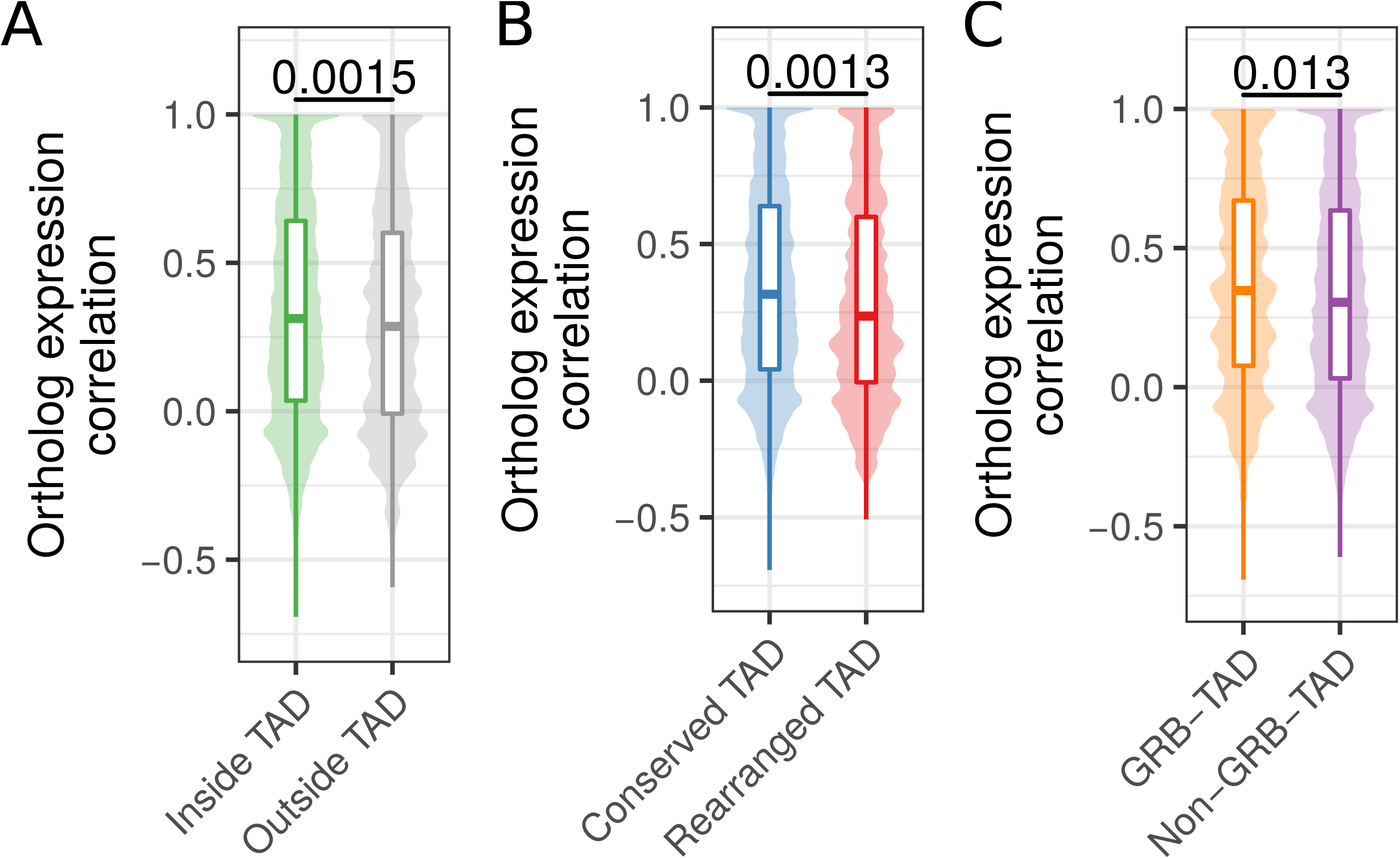
Ortholog gene expression correlation across tissues in conserved and rearranged TADs. **(A)** Expression correlation of orthologs across 19 matching tissues in human and mouse for human genes within or outside of hESC TADs. **(B)** Expression correlation of orthologs across 19 matching tissues in human and mouse for genes in conserved or rearranged TADs. **(C)** Expression correlation of orthologs across 19 matching tissues in human and mouse for genes in GRB-TADs and non-GRB TADs. All P-values according to Wilcoxon rank-sum test.

Next, we further subdivided TADs in two groups, rearranged and conserved, according to syntenic blocks and rearrangements between human and mouse genomes. In brief, a TAD is defined as conserved, if it is completely enclosed by a syntenic alignment block and does not overlap any rearrangement breakpoint. Conversely, a rearranged TAD is not enclosed by a syntenic alignment block and overlaps at least one breakpoint that is farther than 80 kb from its boundary (see Methods). For the hESC TAD data set, this leads to 2,542 conserved and 137 rearranged TADs. The low number of rearranged TADs is consistent with the depletion of rearrangement breakpoints within TADs in general (Fig. 2). In total 8,740 genes in conserved and 645 genes in rearranged TADs could be assigned to a one-to-one ortholog in mouse and are contained in the expression data set. The expression correlation with mouse orthologs were significantly higher for genes in conserved TADs (median R = 0.316) compared to genes in rearranged TADs (median R = 0.237, p = 0.0013) (Fig. 4B). This shows that disruptions of TADs by evolutionary rearrangements are associated with less conserved gene expression profiles across tissues. Although not significant, we also observed a slightly higher expression correlation for 1,003 genes in GRB-TADs compared to 8,038 genes in non-GRB TADs (Fig. 4C, p = 0.13).

In summary, we observed higher expression correlation between orthologs for human genes inside TADs than outside. Moreover, we saw that genes in rearranged TADs show lower gene expression conservation than those in conserved TADs. These results not only support a functional role of TADs in gene regulation, but further support the hypothesis that TAD regions are subjected to purifying selection against their disruption by structural variations such as rearrangements.

## Discussion

Our analysis of rearrangements between human and 12 diverse species shows that TADs are largely stable units of genomes, which are often reshuffled as a whole instead of disrupted by rearrangements. Furthermore, the decreased expression correlation with orthologs in mouse and human in rearranged TADs shows that disruptions of TADs are associated with changes in gene regulation over large evolutionary time scales.

TADs exert their influence on gene expression regulation by determining the set of possible interactions of cis-regulatory sequences with their target promoters [4,6,35]. This might facilitate the cooperation of several sequences that is often needed for the complex spatiotemporal regulation of transcription [36]. The disruption of these enclosed regulatory environments enables the recruitment of other cis-regulatory sequences and might prevent formerly established interactions [37]. The detrimental effects of such events have been shown in the study of diseases [32,38]. There are also incidences where pathogenic phenotypes could be specifically attributed to enhancers establishing contacts to promoters that were formerly out of reach because of intervening TAD boundaries [8,9,39]. This would explain the selective pressure to maintain TAD integrity over large evolutionary distances and why we observe higher gene expression conservation for human genes within TADs compared to genes outside TADs.

Disruptions of TADs by large-scale rearrangements change expression patterns of orthologs across tissues and these changes might be explained by the altered regulatory environment which genes are exposed to after rearrangement [31].

Our results are largely consistent with the reported finding that many TADs correspond to clusters of conserved non-coding elements (GRBs) [26]. We observe a strong depletion of evolutionary rearrangements in GRBs and enrichment at GRB boundaries. This is consistent with comparative genome analysis revealing that GRBs largely overlap with micro-syntenic blocks in *Drosophila* [27] and fish genomes [28]. However, over 60% of human hESC TADs do not overlap GRBs [26], raising the question of whether only a small subset of TADs are conserved. Interestingly, we find also depletion of rearrangements in non-GRB-TADs. This indicates that our rearrangement analysis identifies conservation also for TADs that are not enriched for CNEs. High expression correlation of orthologs in conserved TADs suggestss that the maintenance of expression regulation is important for most genes and probably even more crucial for developmental genes which are frequently found in GRBs.

Previous work using comparative Hi-C analysis in four mammals revealed that insulation of TAD boundaries is robustly conserved at syntenic regions, illustrating this with a few examples of rearrangements between mouse and dog genomes, which were located in both species at TAD boundaries [40]. The results of our analysis of thousands of rearrangements between human and 12 other species confirmed and expanded these earlier observations.

The reliable identification of evolutionary genomic rearrangements is difficult. Especially for non-coding genomic features like TAD boundaries, it is important to use approaches that are unbiased towards coding sequence. Previous studies identified rearrangements by interrupted adjacency of ortholog genes between two organisms [40,41]. However, such an approach assumes equal inter-genic distances, which is violated at TAD boundaries, which have in general higher gene density [3,42]. To avoid this bias we used whole-genome-alignments. However, low quality of the genome assembly of some species might introduce alignment problems and potentially false positive rearrangement breakpoints.

Rearrangements are created by DNA double strand breaks (DSBs), which are not uniquely distributed in the genome. Certain genomic features, such as open chromatin, active transcription and certain histone marks are shown to be enriched at DSBs in somatic translocation sites [43] and evolutionary rearrangements [44–46]. Furthermore, induced DSBs and somatic translocation breakpoints are enriched at chromatin loop anchors [47]. This opens the question of whether our finding of significantly enriched evolutionary rearrangement breakpoints at TAD boundaries could be explained by the molecular properties of the chromatin at TAD boundaries, rather than by the selective pressure to keep TAD function. Although, we cannot distinguish the two explanations entirely, our gene expression analysis indicates stronger conservation of gene expression in conserved TADs and more divergent expression patterns in rearranged TADs. This supports a model in which disruption of TADs are most often disadvantageous for an organism. Structural variations disrupting TADs can lead to miss regulation of neighboring genes as shown for genetic diseases [8,9,32,48] and cancers [49–52].

Interestingly, we observed higher gene expression conservation for human genes within TADs compared to genes outside TADs. The larger syntenic structure of TADs might conserve the regulation likely by maintaining the proximity of promoters and cis-regulatory sequences while genes outside such frameworks are more exposed to changing genomic landscapes, presumably resulting in a greater susceptibility to the recruitment of regulatory sequences.

Apart from the described detrimental effects, our results suggest that TAD rearrangements occurred between genomes of human and mouse and led to changes in expression patterns of many orthologous genes. Since this is likely attributed to changing regulatory environments, it is also conceivable that some rearrangements led to a gain of function. Hence, TAD rearrangements might also provide a vehicle for evolutionary innovation. A single TAD reorganization has the potential to affect the regulation of a whole set of genes in contrast to the more confined consequences of other types of mutations [53]. Since it is also believed that changes in cis-regulatory sequences of developmental genes play a big part in evolutionary innovation [54], the development of the enormous diversity of animal traits in evolution might have been promoted by the rearrangement of structural domains. This is consistent with a model in which new genes can arise by tandem-duplication and during evolution are then re-located to other environments [34]. These changes might have facilitated significant leaps in morphological evolution explaining the emergence of features that could not appear in small gradual steps. Following this hypothesis, TADs would not only constitute structural entities that perform the function of maintaining an enclosed regulatory landscape but could also be a driving force for change by exposing many genes at once to different genomic environments following single events of genomic rearrangement.

## Conclusion

Our results indicate that TADs represent conserved functional building blocks of the genome. We have shown that the majority of evolutionary rearrangements do not affect the integrity of TADs and instead breakpoints are strongly clustered at TAD boundaries. This leads to the conclusion that TADs constitute conserved building blocks of the genome that are often reshuffled as a whole rather than disrupted during evolution. The conservation of TAD regions can be explained by detrimental effects of disrupting cis-regulatory environments that are essential for the spatio-temporal control of gene expression. Indeed we observe a significant association of conserved gene expression in intact TADs and divergent expression patterns in rearranged TADs explaining both why there could be selective pressure on the integrity of TADs over large evolutionary time scales, but also how TAD rearrangement can explain evolutionary leaps.

## Methods

### Rearrangement breakpoints from whole-genome alignments

Rearrangement breakpoints were identified between human and 12 selected vertebrate species from whole-genome-alignment data (Table 1). Alignment data were downloaded as net files from UCSC Genome Browser for human genome hg38 and the genomes listed in Table 1. The whole-genome data consists of consecutive alignment blocks that are chained and hierarchically ordered in the so-called nets [19]. Chains represent blocks of interrupted syntenic regions and may include larger gaps. When hierarchically arranged in a net file, child chains can complement their parents when they align nearby segments that fill the alignment gaps of their parents but may also break the synteny when incorporating distal segments. We implemented a computer program to extract rearrangement breakpoints from net files based on the length and type of fills. Start and end points of top-level or non-syntenic fills are reported as rearrangement breakpoint if the fill exceeds a given size threshold. We used different size thresholds to optimize both the number of identified breakpoints and to avoid biases of transposable elements that might be responsible for many small interruptions of alignment chains. In this way, we extracted rearrangement breakpoints between human and 12 genomes using size thresholds of 10 kb, 100 kb, and 1000 kb. To compare breakpoints to TADs we converted the breakpoint coordinates from hg38 to hg19 genome assembly using the liftOver tool from UCSC Genome Browser [55].

**Table 1.** Species used for breakpoint identification from whole-genome alignments with human.

| Common name | Species | Genome Assembly | Divergence to human (mya) |
| --- | --- | --- | --- |
| Chimpanzee | *Pan troglodytes* | panTro5 | 6.65 |
| Gorilla | *Gorilla gorilla gorilla* | gorGor5 | 9.06 |
| Orangutan | *Pongo abelii* | ponAbe2 | 15.76 |
| Rhesus | *Macaca mulatta* | rheMac8 | 29.44 |
| Mouse lemur | *Microcebus murinus* | micMur2 | 74 |
| Mouse | *Mus musculus* | mm10 | 90 |
| Cattle | *Bos taurus* | bosTau8 | 96 |
| Manatee | *Trichechus manatus latirostris* | triMan1 | 105 |
| Opossum | *Monodelphis domestica* | monDom5 | 159 |
| Chicken | *Gallus gallus* | galGal5 | 312 |
| Clawed frog | *Xenopus tropicalis* | xenTro7 | 352 |
| Zebrafish | *Danio rerio* | danRer10 | 435 |

### Topologically associating domains and contact domains

We obtained topologically associating domain (TAD) calls from published Hi-C experiments in human embryonic stem cells (hESC) [3] and contact domains from published *in situ* Hi-C experiments in human GM12878 cells [22]. Genomic coordinates of hESC TADs were converted from hg18 to hg19 genome assembly using the UCSC liftOver tool [55].

### Breakpoint distributions at TADs

To quantify the number of breakpoints around TADs and TAD boundaries we enlarged TAD regions by 50% of their total length on each side. The range was then subdivided into 20 equal sized bins and the number of overlapping breakpoints computed. This results in a matrix in which rows represent individual TADs and columns represent bins along TAD regions. The sum of each column indicates the number of breakpoints for corresponding bins and therefore the same relative location around TADs. For comparable visualization between different data sets, the column-wise summed breakpoint counts were further normalized as percent values of the total breakpoint number in the matrix.

### Quantification of breakpoint enrichment

To quantify the enrichment of breakpoints at domain boundaries, we generated random breakpoints as background control. For each chromosome, we placed the same number of actual breakpoints at a random position of the chromosome. For each breakpoint data set we simulated 100 times the same number of random breakpoints. We then computed the distribution of random breakpoints around TADs in the same way as described above for actual breakpoints. To compute enrichment of actual breakpoints compared to simulated controls, we classified each breakpoint located in a window of 400 kb around TAD borders in either close to a TAD boundary, if distance between breakpoint and TAD boundary was smaller or equal to 40 kb or as distant, when distance was larger than 40 kb. This results in a contingency table of actual and random breakpoints that are either close or distal to TAD boundaries. We computed log odds ratios as effect size of enrichment and p-values according to Fishers two-sided exact test. Additionally, we compared the distance of all actual and random breakpoints to their nearest TAD boundary using the Wilcoxon’s rank-sum test.

### Expression data for mouse and human orthologs

Promoter based expression data from CAGE analysis in human and mouse tissues from the FANTOM5 project [33] were retrieved from the EBI Expression Atlas [55] as baseline expression values per gene and tissue. The meta data of samples contains tissue annotations as term IDs from Uberon, an integrated cross-species ontology covering anatomical structures in animals [56]. Human and mouse samples were assigned to each other if they had the same developmental stage and matching Uberon term IDs. This resulted in 19 samples for each organism with corresponding tissues.

We used the R package biomaRt to retrieve all human genes in the Ensembl database (version grch37.ensembl.org) and could assign 13,065 to ortholog genes in mouse by allowing only the one-to-one orthology type [56]. Of these ortholog pairs, 12,696 are contained in the expression data described above. For each pair of orthologs we computed the correlation of expression values across matching tissues as Pearson's correlation coefficient.

### Classification of TADs and genes according to rearrangements and GRBs

We classified hESC TADs according to rearrangements between human and mouse genomes. We define a TAD as conserved if it is completely enclosed within a fill in the net file and no rearrangement breakpoint from any size threshold is located in the TAD region with a distance larger than 80 kb from the TAD boundary. A TAD is defined as rearranged, if the TAD is not enclosed completely by any fill in the net file, overlaps at least one breakpoint inferred using a 1000 kb fill size threshold, and this breakpoint is further than 80 kb away from each TAD boundary. TADs were also classified according to their overlap with GRBs as in [26]. A given TAD is a GRB-TAD if it overlaps with more than 80% of the TAD size with a GRB. A TAD is classified as non-GRB if it has less than 20% overlap with GRBs. The 12,696 human genes with mouse ortholog and expression data were grouped according to their location with respect to hESC TADs. We used the transcription start site (TSS) of the longest transcript per gene to group each gene as within TAD if the TSS overlaps a hESC TAD or as outside TADs, if not. Furthermore, we grouped genes in TADs according to conserved or rearranged TADs and separately according to GRB and non-GRB TADs.

### Source code and implementation details

The source code of the entire analysis described here is available on GitHub: https://github.com/Juppen/TAD-Evolution. The identification of breakpoints and extraction of fills from whole-genome alignment data was implemented in Python scripts. Reading of BED files and overlap calculations with TADs and TAD bins were computed in R with Bioconductor [57] packages `rtracklayer` [58] and `GenomicRanges` [59]. Gene coordinates and ortholog assignments were retrieved from Ensemble data base (version grch37.ensembl.org) using the package `biomaRt` [60]. For data integration and visualization we used R packages from `tidyverse` [61].

## Declarations

### Ethics approval and consent to participate

Not applicable

### Consent for publication

Not applicable

### Availability of data and material

The source code of all analysis is available on GitHub: https://github.com/Juppen/TAD-Evolutio. All the genomic data used for analyses are freely available to be downloaded from the UCSC Genome Browser and EBI Expression Atlas with identifiers listed in Table 1 and S1.

### Authors’ contributions

JK and JI developed and implemented the methods and performed the analysis. JI conceived the study. JK wrote the first draft of the manuscript. JK, MA and JI wrote the manuscript. MA supervised the study.

## Acknowledgments

The authors thank all members of the CBDM group for fruitful discussions.

## Supplementary Data

**Table S1**

**Matching tissues and samples with CAGE expression data in human and mouse.**

**Table S2**

**Ortholog genes in human and mouse with gene expression correlation across tissues.**

**Figure S1**

**Distribution of evolutionary rearrangement breakpoints between human and 12 vertebrate genomes around domains.** Relative breakpoint numbers from human and different species (horizontal panels) around hESC TADs (left), GM12878 contact domains (center), and GRBs (left). Blue color scale represents breakpoints from different fill-size thresholds. Dotted lines in gray show simulated background controls of randomly placed breakpoints.

**Figure S2**

**Distance between rearrangement breakpoints and random controls to closest TAD boundary**. For each species (y-axis) and fill size threshold (vertical panels) the distances from all identified rearrangement breakpoints to its closest TAD boundary (x-axis) are compared between actual rearrangements (blue) and 100 times randomized background controls (gray). The left panel shows distances to next hESC TAD boundary and the right panel distances to closest GM12878 contact domain boundary. P-values according to Wilcoxon's rank-sum test.

